# Sequences Dimensionality-Reduction by *K*-mer Substring Space Sampling Enables Effective Resemblance- and Containment-Analysis for Large-Scale omics-data

**DOI:** 10.1101/729665

**Authors:** Huiguang Yi, Yanling Lin, Wenfei Jin

## Abstract

We proposed a new sequence sketching technique named *k*-mer substring space decomposition (kssd), which sketches sequences via *k*-mer substring space sampling instead of local-sensitive hashing. Kssd is more accurate and faster for resemblance estimation than other sketching methods developed so far. Notably, kssd is robust even when two sequences are of very different sizes. For containment analysis, kssd slightly outperformed mash screen—its closest competitor—in accuracy, while took testing datasets of 110,535 times less space occupation and consumed 2,523 times less CPU time than mash screen—suggesting kssd is suite for quick containment analysis for almost the entire omics datasets deposited in NCBI. We detailed the kssd algorithm, provided proofs of its statistical properties and discussed the roots of its superiority, limitations and future directions. Kssd is freely available under an Apache License, Version 2.0 (https://github.com/yhg926/public_kssd)

## 1 Introduction

Up to this manuscript is composing, the size of NCBI SRA database alone has reached to 30 peta base-pairs, the number is seemly will grow exponentially in the near future [1]. Additionally, the continuously emerging independent projects such as The Cancer Genome Atlas [2], 1000 Genomes [3], Human Microbiome Project [4], The Genotype-Tissue Expression (GTEx) project [5], Human Cell Atlas [6], etc. produced so-called omics-datasets (including genomics, metagenomics, transcriptomics, etc.) of terabytes to petabytes each [7]. Such a phenomena of omics-data explosion bring us to the ‘omics big data’ era [8]. On the one hand, the ‘omics big data’ is revolutionizing biomedical science profoundly [9]; on the other, such volume of data put a heavy burden for data storage and operations, and put a pressing need for effective sequence dimensionality-reduction techniques.

Recently, the omics ‘big data’ problem was largely alleviated by a class of sequence dimensionality-reduction techniques—minhash sketching, including methods such as mash [10], bindash [11], etc. Minhash sketching selects fix number (typically around 1000) of *k*-mers, termed sketch, which have minimum hashes among all the *k*-mers from the sequence, for the reduced representation of this sequence. No matter how large the original sequences are, taking their sketches of merely several Kbytes in size, minhash sketching could effectively estimate the pairwise distances, hence enabled first time rapid clustering of entire NCBI RefSeq. Other applications of minhash sketching include identifying mis-labeled samples, real-time outbreak pathogen surveillance, searching genomics database, etc. [10]. Notably, minhash sketching also play a crucial part in single-molecular long-reads assembly by overcome the computational bottleneck of all vs. all long-reads overlapping [12].

However, minhash sketching is not suite for comparing two *k*-mer sets of very different sizes [13], which is the scenario of many potential applications, e.g. prioritizing the reference genomes for a metagenomics dataset, composition analysis of a metagenomics dataset, searching viruses/bacteria contaminations in human omics samples, etc. Mash screen (mash version 2) then was developed to address this problem via estimating the proportion (termed containment-index or containment-coefficient) of a reference (the smaller *k*-mer set) contained in a sequences-mixture (the larger *k*-mer set). The containment-coefficient then is converted to containment-score representing identity between the two *k*-mer sets (see [14] for the definition); if there are many mixtures/references to be compared, the references are sketched as the reference-sketches database, then the mixtures are queried one by one against the reference-sketches database to calculate the containment-coefficient of each reference in each mixture [14]. But the mixtures, which consume the majority of the space, are not sketched at all, hence the merits of sketching technique—extremely high space- and time-efficiency—are largely lost, which make mash screen more suitable for answering the question “what reference genomes contained in my(one) omics run?” [14], but not “what reference genomes contained in which omics run?”.

To address the latter question, we here propose a new sequence sketching technique named *k*-mer substring space decomposition (kssd) which use random *k*-mer substring space sampling (see Methods) instead of minhash or more generally local-sensitive hashing [15] for sequences sketching. Kssd sketches both sequences-mixtures and references and can effectively estimate the distances using sketches even if they are of very different sizes. Conventionally, sketching technique is used to measure the extents of the two kinds of relationships— resemblance and containment, which capture the notions of “roughly the same” and “roughly contained” for two sequences (or documents), respectively [13]; more specifically, the resemblance captures the distance/similarity of two sequence of similar sizes, whereas the containment captures the distance/similarity of two sequence of very different sizes. Here we examine the performances of kssd and minhash-based sketching methods w.r.t. both resemblance- and containment-estimation.

## 2 Results

### 2.1 Accuracy of resemblance estimation

To compare the accuracy of resemblance estimation of kssd with other sketching methods, we evolved a reference genome in-silico with 600 pre-defined mutation rates range from 0.001 to 0.60 with stepwise increase of 0.001 (use an in-house script, see supplementary file 1 for the dataset) which were served as ground truths hereafter. The mutants were divided into closely-related group and distantly-related group (with mutation rate 0. 001 ~ 0.3 and 0.301 ~ 0.60, respectively). The mutation distances between the reference and its mutants were estimated then by kssd and two other sketching methods mash and bindash, with variated *k*-mer lengths and sketch-sizes. The Pearson correlation-coefficient of the ground truths and the estimates was adopted as the accuracy-measurement (Figure 1A, B).

**Figure 1.**
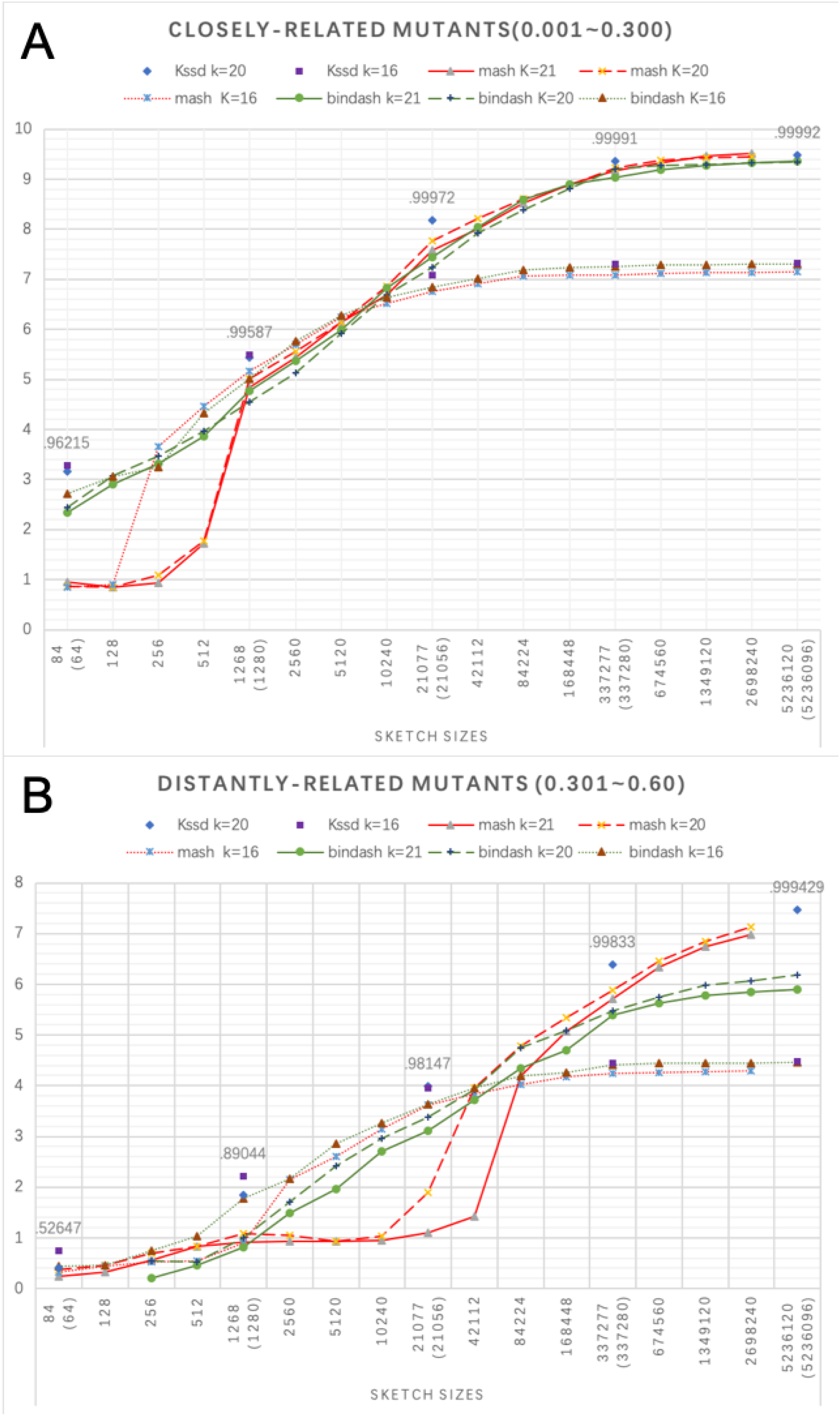
Accuracy of kssd, mash and bindash. (A) Pearson correlation-coefficients on closely-related group and (B) distantly-related group. y-axis is scaled to −**log**(**1** — ***r***) for plotting clarity, where *r* is the Pearson correlation-coefficient between the ground truth- and the estimated-mutation rates. The decimal above the highest data point at a sketch-size is the maximal *r* value of all the three methods of all *k* settings with that sketch-size. The default *k*-mer lengths *k* for kssd, bindash and mash are 16, 21 and 21, respectively; to match *k*, we also run mash and bindash with *k* = 16 in addition to the default *k* settings, but kssd takes only even *k*, so we also run with *k* = 20 to variate *k*. Due to different sketching mechanism, mash and bindash take as parameter the sketch-size of continuous integers and multiples of 64, respectively; but sketch-size is not a parameter of kssd and can only counted from the sketch file. To match sketch-sizes as closely as possible, we first sketched the reference using kssd with dimensionality-reduction levels *n* = {4, 3, 2, 1, 0} and obtained the sketch-sizes *s_k_* = {84, 1268, 21077, 337277, 5236120}, respectively; we got the nearest multiples of 64 of *s_k_* (the parenthesized values) and interpolated with their 2-, 4- and 8-fold sketch-sizes to obtain the sketch-sizes parameter *s_b_* for bindash; and we merged *s_k_* and the interpolated points of *s_b_* to obtain the sketch-size parameter *s_m_* for mash. Mash with *k*=20, 21 at sketch-size 5236120 is not shown due to the running error. Bindash with *k*=20, 21 at sketch-size 64 and 128 on distantly-related group are not shown due to the estimates lacking of variation for correlation.

The results show clear stratifications w.r.t. *k*-settings, where close *k* (20 vs. 21) yield generally similar curves whereas distant *k* (16 vs. 20, 21) yield dissimilar curves. When the sketch-size is relative smaller, using shorter *k* (16 here) is more accurate than longer *k* (20, 21 here); but when the sketch-size is larger (larger than 10,240 and 42,112 in closely-related group and distantly-related group, respectively), using longer *k* is more accurate (Figure 1A, B). However, for the sake of efficiency, sketching technique rarely use sketch-size greater than 10,000, so the shorter *k* is preferred. Given the same *k*-settings (*k* =16 or 20), kssd outperforms both mash and bindash on both groups, especially when the sketch-size is small (84, 1,268 and 21,077), where the accuracy of kssd is on par with that of mash/bindash with doubled sketch-size (Figure 1).

### 2.2 Accuracy of containment estimation

To compare kssd with mash screen for the accuracy of containment estimation, we benchmarked kssd by the same approach used by Ondov et al. [14], where a constitutes-known synthetic microbiome community (referred as the shakya dataset, see supplemental file 2 for the 64 known constitutes), was adopted as the testing dataset [14]. Notably, due to sample contaminations, the real shakya dataset (accession: SRR606249) contains also exogenous genomes that are not included in the 64 known constitutes [16], and the contaminations could be revealed by comparing to the simulated shakya dataset that consisted of only the short-reads from the 64 known constitutes (served as the contamination-free control, see simulation method in [14]). However, Ondov et al. did not provide the exact list of the references they used, so we took as references the latest NCBI prokaryotic assembly which consisted of 138,743 genomes (see supplement file 2 for the summary) instead of the 82,086 genomes of Ondov et al.

Given the references, the containment-measurements (namely, the aaf-distance of kssd and the containment-score of mash screen, see Equation 5 and [14] for the respective definitions) of each reference to the real- and the simulated-shakya datasets were plotted against the minimum of the distances of this reference to the 64 known constitutes (referred as the minimum reference-to-constitutes distance hereafter, Figure 2). The Fig. 3 in the paper of Ondov et al. [14] were faithfully reproduced (Figure 2, 2^nd^ row), though we used more references here. The kssd containment-analysis (Figure 2, 1^st^ row) conveys almost the same information with that of mash screen, where the data points are biased due to the same low-abundance constitutes (Figure 2, circled in purple) and the same contaminations (Figure 2, circled in orange and red). However, the data points for mash screen get increasingly discrete with the increase of the minimum reference-to-constitutes distance (*x*-axis) or the decrease of containment-score (*y*-axis) due to its sketch-size is fixed. In contrast, the data points for kssd are continuous with the change of *x* and *y*, which is more realistic. Since kssd estimates the distance (Equation 5) whereas mash screen estimates the identity between the reference and the shakya dataset, the containment-measurement of kssd is positively correlated with the minimum reference-to-constitutes distance whereas that of mash screen is negatively correlated. Therefore, we used the absolute correlation-coefficients for accuracy comparison. Both the kssd and the mash screen containment-measurements are correlated well with the minimum reference-to-constitutes distances less than 0.15, but kssd slightly outperforms mash screen, where the absolute correlation-coefficients of kssd for the simulated- and the real-shakya dataset are 0.9869 and 0.9611, respectively, and of mash screen are 0.9808 and 0.9592, respectively.

**Figure 2.**
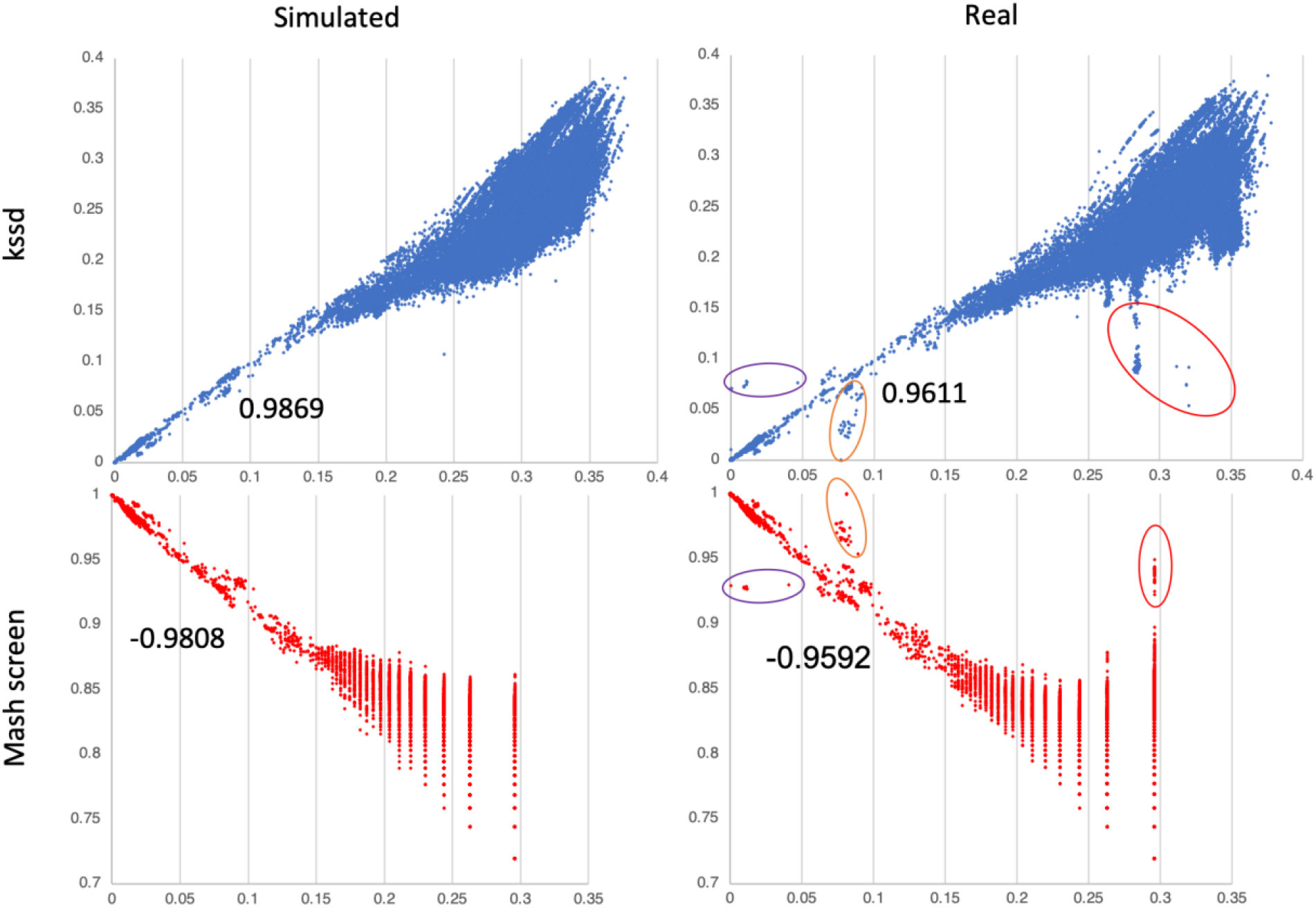
accuracy of containment-measurements. *x*-axis indicates the minimum reference-to-constitutes distance computed by kssd (1^st^ row or blue) or mash (2^nd^ row or red), and *y*-axis indicates the reference-to-mixture containment-measurement (aaf-distance or containment-score for kssd or mash screen, respectively). The decimals below the data points are the correlation-coefficients of the plot controlling *x* < 0.15. The 1^st^ and 2^nd^ columns are the plots of the simulated and the real shakya datasets, respectively. On the real shakya datasets, the data points biased from expect due to low abundance constitutes are circled in purple, and those biased due to two different contaminations are circled in orange and red.

Notably, kssd took as queries the sketches of the shakya datasets combined of only 580Kbytes—110,535 times less space occupation than that of mash screen which took the raw datasets combined of 52Gbytes. Moreover, kssd took only 23.5 CPU minutes for the raw dataset sketching and 3 CPU seconds for the containment estimation, whereas mash screen took totally 126 CPU minutes for screening and the containment estimation. There is no need for kssd to sketch the raw datasets again for the containment-analysis for new references whereas mash screen would take another 126 CPU minutes for another 138,743 new references, which is likely happen in the near future since the NCBI Refseq database is growing rapidly [17], meaning the asymptotic time-consumption of kssd is only 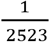 of that of mash screen (3 CPU seconds vs. 126 CPU minutes). Such a high space- and time-efficiency of kssd should enable building a sketch database for almost all the short-read datasets deposited in NCBI for fast containment-analysis.

### 2.3 Computational efficiency

We also tested the time-complexities of kssd, mash and bindash using a dataset consisted of 100K bacteria reference genomes under a 12-cores machine. It took 10531, 19094 and 328835 CPU seconds for kssd, bindash and Mash, respectively and took 933, 11,335 and 27,600 elapsed seconds for kssd, bindash and mash, respectively (Figure 3A and B) to complete the all pairwise distance computation. Therefore, kssd is 12 times and 30 times faster than bindash and mash respectively while using same number of threads (12 threads here). The comparison of time-complexity was conducted using similar sketch-size (1,780 and 2,048 for kssd/mash and bindash, respectively), as illustrated in Figure 1, kssd can achieved similar accuracies by using only half of the sketch-sizes of mash/bindash. Therefore, kssd could outperform mash/bindash even more under the accuracy criterion.

**Figure 3.**
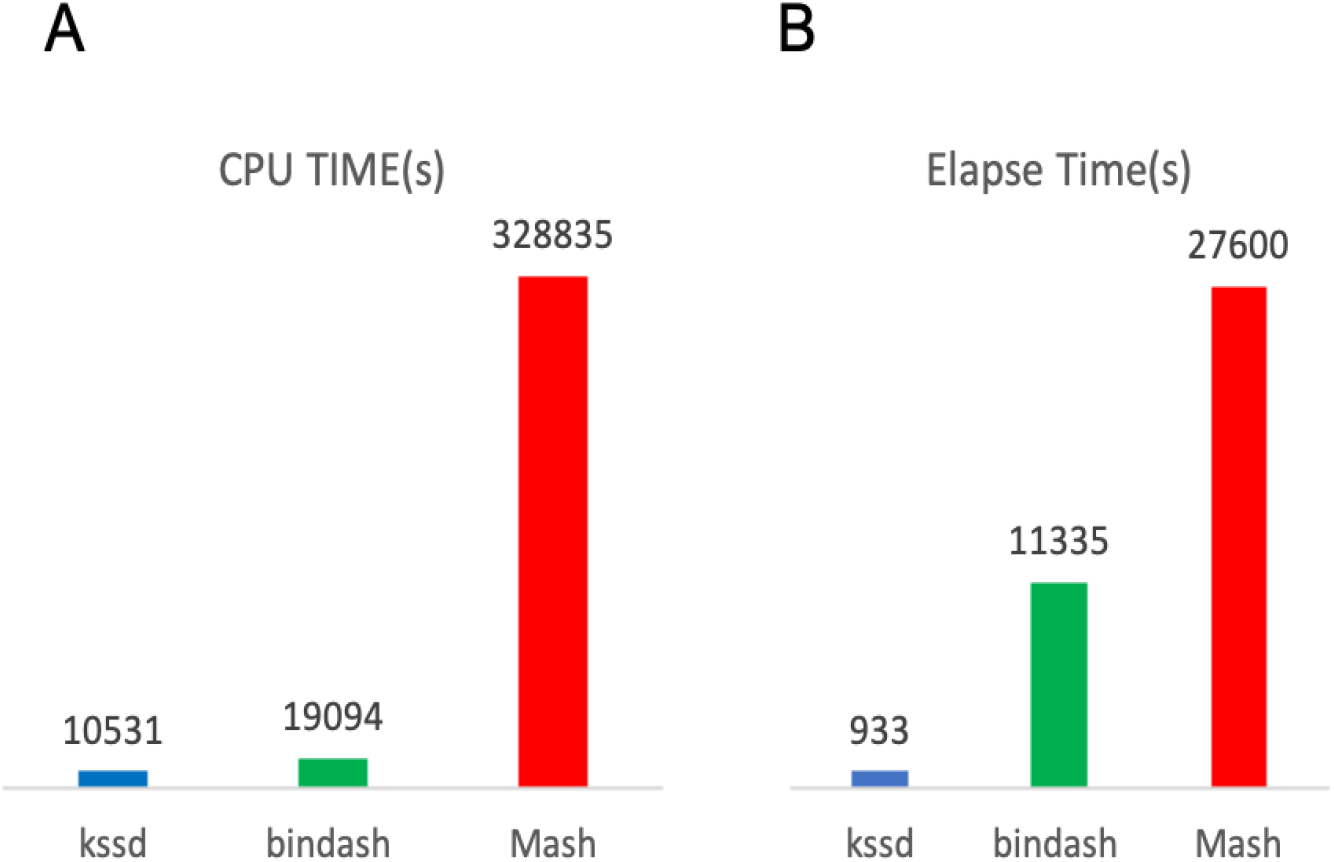
Computational efficiencies. Assessments of CPU time (A), Elapse time (B) of the three methods were performed on a 32Gb,12-cores machine using a test dataset consisting of 100k bacteria genomes.

## 3 Discussion

We showed in this manuscript that kssd has greater accuracy and computational efficiency versus minhash sketching methods, and notably, kssd distance estimation is robust even when two sequences are of very different sizes. Mash screen estimates containment-coefficients from the raw data of the sequences-mixtures, whereas kssd estimates from the sketches which are hundreds of thousands of times smaller. Such a high-efficient summarizing ability of kssd facilitates large-scale containment analysis. We illuminated that the Jaccard- and the containment-coefficients estimated by kssd are essentially the sample proportions which are asymptotically gaussian distributed. Therefore, the standard deviations and 95% CIs could be resolved analytically (see Methods), which is not yet framed by other sketching methods. Kssd is also the only method able to decompose a *k*-mer set into a collection of mutual-exclusive sketches, which enables calculating the ground truth Jaccard- and containment-coefficient even when fully loading of the whole *k*-mer set is restrained by the memory limitation.

Kssd first selects *k*-mers and then maps the selected *k*-mers to integers one-to-one using a pre-determined recoding scheme. This process is reversible and could be seen as a special hash function that has no hash-collision (namely, two different *k*-mers are mapped to the same integer). In contrast, other sketching methods typically first hash from the whole *k*-mers space to a much smaller integers space, which caused hash-collision, before the hashes are selected for sketching. Since the final sketch would be smaller than the integers space, sketching methods like kssd putting *k*-mers selection before hashing could avoid hash-collision. Hash-collision, usually infrequent though, would increase the variation of the Jaccard- or containment-coefficient estimates. Thanks to the resolution of the hash-collision, kssd acquires its superior accuracy over other methods.

Though kssd possess the above-mentioned merits, there are still limitations: Firstly, Kssd loses its efficiency when searching individual-gene- or small-virus-references contained in sequences-mixtures, since these sequences are too short to be dimensionality-reduced further to an informative sketch [14]. And kssd requires the sequences-mixtures to be dimensionality-reduced with the same rate with the references, so the sequences-mixtures cannot be dimensionality-reduced either. We suggest the references should be at least 10K base-pairs, so that the lowest-level sketches (with 16 folds of dimensionality-reduction) would still be informative enough for effective distance estimation. For single-gene searching, the tool BIGSI [18] would be a more suitable choice, and for numerous small-virus genomes searching, mash screen would be better. Secondly, in current version of kssd implementation, the choices for the dimensionality-reduction rate are restricted to 16^*n*^ (*n* ranges from 0 to 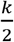, where *k* is *k*-mer length and kssd only work with even *k*), which prevents fine-tuning of the dimensionality-reduction rate. In addition, kssd currently cannot handling amino-acid sequences and 10X genomic single-cell sequences, and cannot overlapping single-molecular-long-reads (the latter two require sketching multiple sequences in a single file individually). Extensions including these functionalities are our future directions.

Another important direction is to estimate directly from the sketch the abundance of each reference contained. Though it is easy to tracking in sketch the *k*-mer occurrences from the sequences-mixture, the relationship between the *k*-mer occurrences and the abundance of each reference contained is complicated, since other genomes contained may also contribute to the *k*-mer occurrences thus confounding abundance estimation. However, the relationship is not totally chaotic, it is evidenced that there is clear average nucleotide identity (ANI) boundary (ANI = 95%) between two difference prokaryotic species [19], which means roughly 5% or larger genome regions are species-specific, thus the *k*-mer occurrences of these regions are not or much less confounded by other species and probably reflect the truth abundance. Therefore, a robust inferring of species-specific *k*-mer occurrences would be the key for sketch-based abundance estimation.

Though minhash sketching was introduced for omics-data handling recently, it has long history of development that initially used for detecting similar web pages and clustering documents [13]. In contrary, kssd was introduced first time in this study, but we speculate that a modified version of kssd can perform tasks like documents clustering or similar web pages detection as well. After all, the documents, webpages content and DNA are essentially all sequences but with a different alphabet set.

## 4 Methods

### 4.1 The main idea

The kssd idea originated from the naive sketching method of sampling *k*-mers directly from the sequence as its sketch. However, such a sketching method is ill-suited for distance estimation since the two sketches *S*(*A*) and *S*(*B*) are probably drawn from two unrelated regions of two genome *A* and *B*. Therefore, they have very few shared *k*-mers (with a Jaccard-coefficient approximate to 0), even when *A* and *B* are nearly identical (Figure 4A). Notwithstanding its naivety, this thought inspired us the idea of *k*-mer space sampling—Firstly, a subset of *k*-mers *s*, termed *k*-mer subspace, is drawn randomly from *k*-mer space ***S*** (namely the collection of all possible string of length *k* defined in a given alphabet set Σ); then the sketch of any given sequence is built by overlapping *s* with the *k*-mers set of this sequence. Since *s* is an unbiased sampling of the *k*-mer space ***S***, it is independent of any instance *k*-mer sets. After sketching, two sequences *A* and *B*, their intersection *A* ∩ *B* and union *A*∪*B* should go through dimensionality-reductions of the same expectation rate of 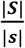. Therefore, it enables measuring both the resemblance and the containment of the two sequences directly using their sketches, even if they are of very difference sizes (Figure 4B). The *k*-mer space sampling could be extended to *k*-mer space shuffling—where *k*-mer space ***S*** is shuffled and split into *N* subspaces of equal size, so that *N* sketches could be obtained for a *k*-mer set by sketching using the *N* subspaces individually. All the *N* sketches combined is a lossless representative of the original *k*-mer set, such a lossless property is particularly useful when the ground truth Jaccard- or containment-coefficient is needed when the memory is limited.

**Figure 4.**
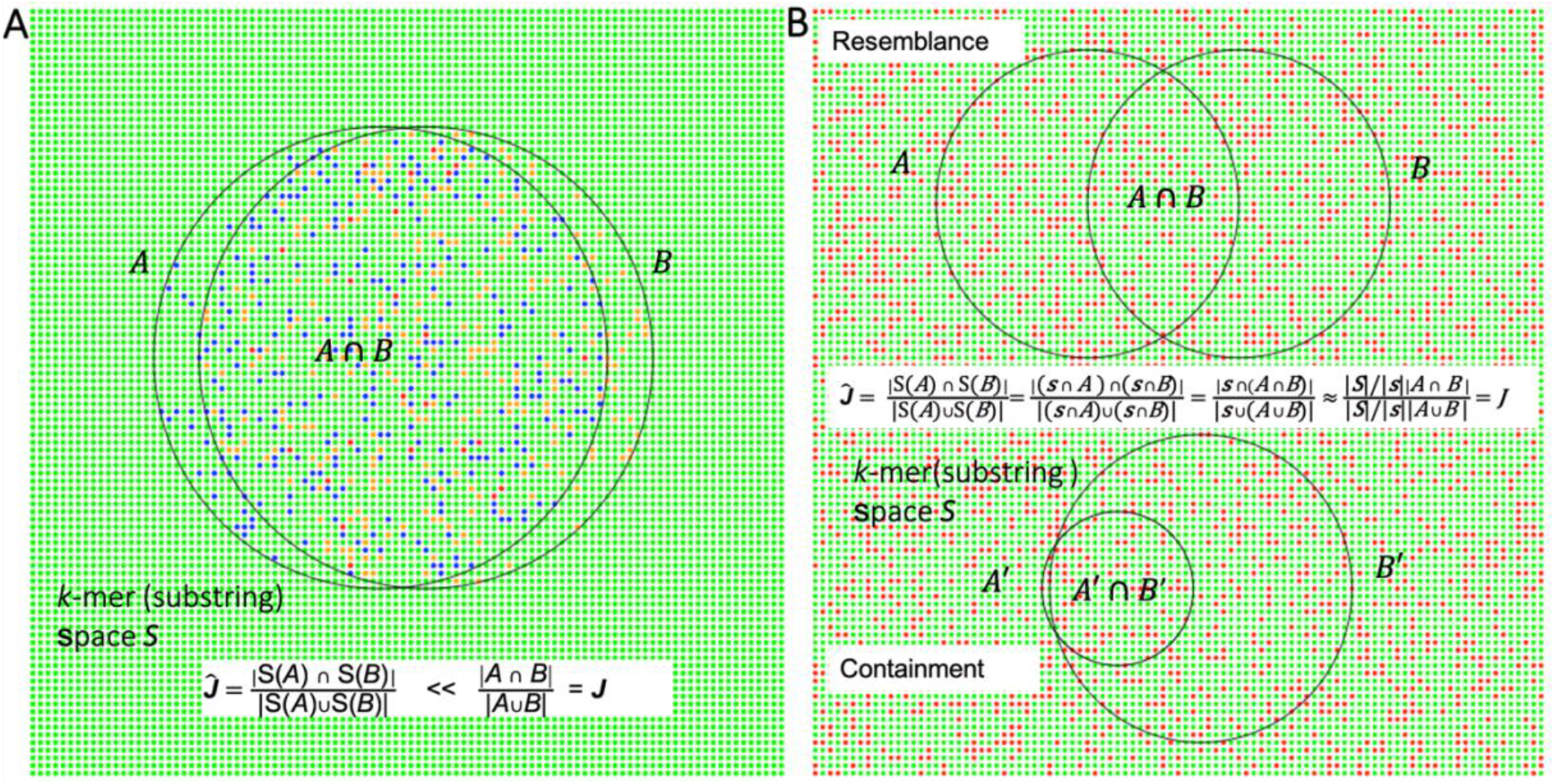
The main idea of kssd. (A) The set of blue dots *S*(*A*) and the set of orange dots *S*(*B*) are the *k*-mer sets randomly drawn from sequence *A* and *B*, respectively, and the set of red dots is *S*(*A*) ∩ *S*(*B*). (B) The set of all red dots is the *k*-mer subspace *s* that sampling from the *k*-mer space ***S***, where ***S*** = green dots ∪ red dots.

However, *k*-mer space sampling/shuffling is computational expensive when *k* is large, since it consumes O(|Σ|^*k*^) memory and time. For example, when *k* = 22, a common *k* choice for mammalian genome analysis, it occupies 16Tbytes memory and takes 4^22^ CPU operations (|Σ| = 4 here) for the *k*-mer space shuffling alone, which is an unrealistic resource demand. To address this, we generalized *k*-mer space sampling/shuffling to *k*-mer substring space sampling/shuffling (kssd)—where a substring of the *k*-mer is selected according to a predefined pattern, so that we can use *k*-mer substring space sampling/shuffling instead of whole *k*-mer space sampling/shuffling (Figure 5). In this way, the computational complexity is dramatically reduced.

**Figure 5.**
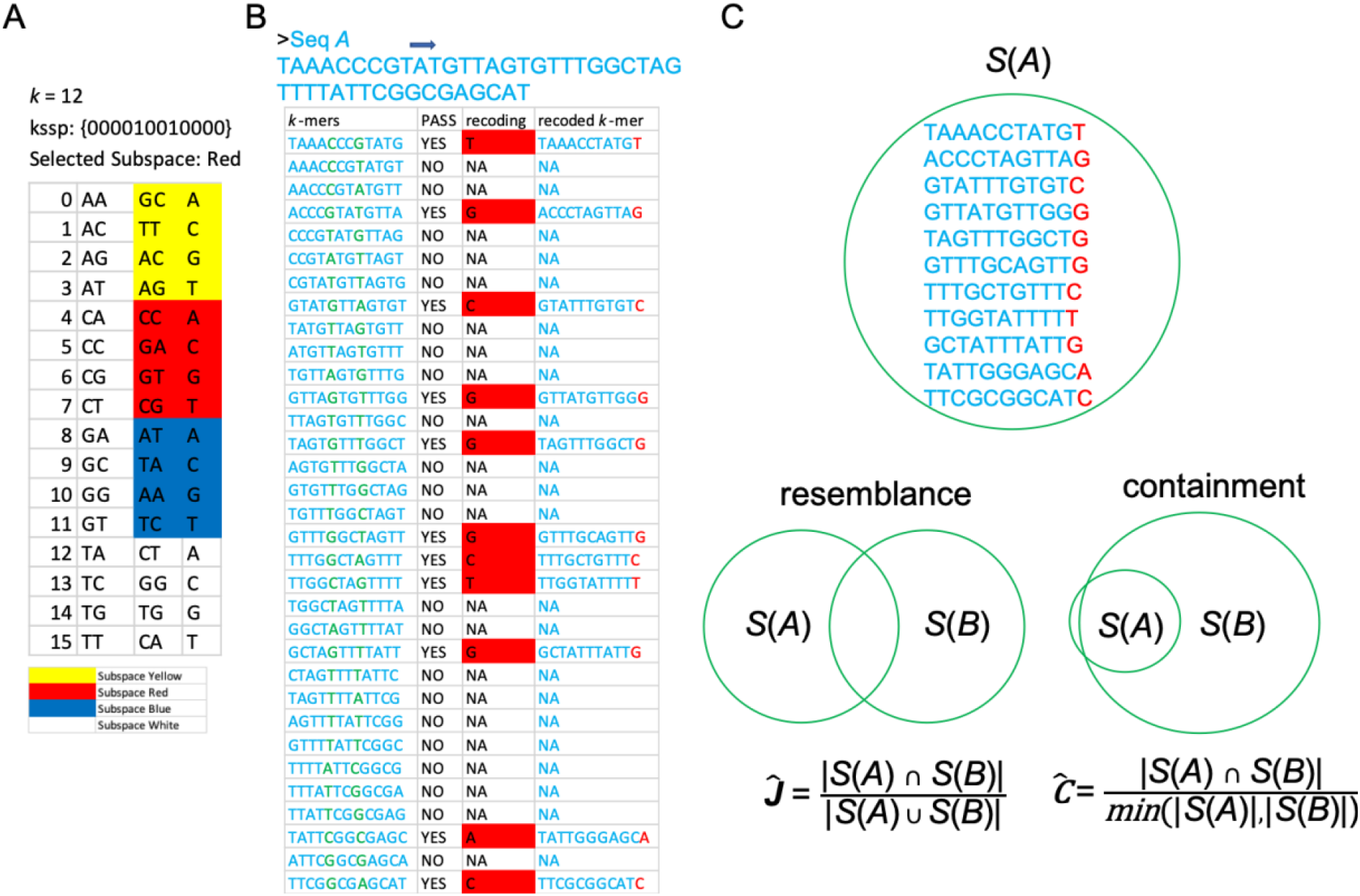
kssd algorithm overview. (A) *k*-mer substring space shuffling. First, a *k*-mer substring selection pattern (kssp) *p* = ‘000010010000’ is pre-determined for the 12-mer analysis, so the length of *p*-selected-substring is 2 and the *k*-mer substring space has dimensionality *D* = 4^2^ =16 (|***Σ***| = 4 for nucleotide sequence, 1^st^ and 2^nd^ column). This 16-dimensions space is shuffled and partitioned into *N* subspaces of equal size (3^rd^ column, *N* = 4 here), and the dimensions in each subspace are recoded by the lexically-ordered strings of length 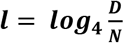 (4^th^ column, ***l*** =1 here). We use one subspace ***s*** (3^rd^ and 4^th^ column, the red subspace here) for sequence sketching. (B) Sequence sketching. First the *k*-mers with *p*-selected-substring (green substring in 1^st^ column) belonging to the red subspace *s* are selected (2^nd^ column), where the *p*-selected-substrings are recoded by the lexically-ordered dimension (3^rd^ column), thus each of the selected *k*-mers is rearranged in such a way that the recoded *p*-selected-substring suffixes the rest substring (4^th^ column). (C) Kssd distance. Once all the sequences are sketched, the Jaccard- and containment-coefficients could be estimated by 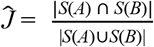 and 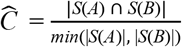, respectively.

### 4.2 Mathematical characterization of kssd method

#### 4.2.1 *k*-mer substring selection pattern, *k*-mer substring space sampling/shuffling and sketching

A *k*-mer substring selection pattern (kssp) *p* is a ‘01’ string of length *k*, by which the letters at the 1s of the given *k*-mer (or *k*-mer set) *K* are concatenated, termed as the *p*-selected-*K*-substring(s) and denoted by *p*^1^(*K*), and the letters at 0s are also concatenated, termed as *p*-unselected-*K*-substring(s) and denoted by *p*^0^(*K*). The weight of *p*, denotated by *w*, is the number of 1s in *p* (also the length of *p*^1^(*K*)). Given alphabet set *Σ* and weight *w*, the *k*-mer substring space ***S*** is the collection of all length *w* strings defined in *Σ*, thus has dimensionality |***S***| = |*Σ*|^*W*^ (we view each element in ***S*** as a dimension). ***S*** is first Fisher-Yates shuffled [20] and then partitioned into *N* subspaces of equal size, denoted by ***s_1_, s_2_***, …, ***s_N_***; for computational convenience, we choose *N* = |*Σ*|^*m*^, where *m* could be any positive integer ≤ *w*, so that |*s*| = |*Σ*|^*w-m*^ (***s*** ∈ { ***s_1_, s_2_*** …, ***s_N_*** }), and the dimensions of *s* could be represented by lexically-ordered strings of length *w-m* (Figure 5A).

Let *A* be a *k*-mer set, the subset *R*(*A*) = {*K*|*K* ∈ *A, p*^1^(*K*) ∈ ***s***} is mapped to sketch *S*(*A*) as follow: for each *k*-mer *K* ∈ *R*(*A*), the *p*-selected-*K*-substring *p*^1^(*K*) is recoded by the lexical order of its dimension in the subspace ***s***, denoted by *s*(*p*^1^(*K*)), then the *p*-unselected-*K*-substring *p*^0^(*K*) and the string *s*(*p*^1^(*K*)) are concatenated into the new string *p*^0^(*K*) *s*(*p*^1^(*K*)), which has length of *k-m* (Figure 5B). This *k*-mer recoding scheme could be summarized by the function *r*(*K*) = *p*^0^(*K*)*s*(*p*^1^(*K*)), and the overall sketching process could be summarized by:

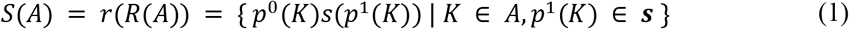

#### 4.2.2 kssd distance

The Jaccard- and the containment-coefficients for a *k*-mer sets pair (*A, B*) are estimated by:

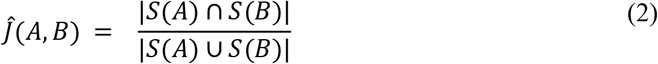

and

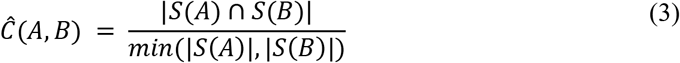

respectively. *Ĵ* and *Ĉ* could be further converted to mash- and aaf-distances by

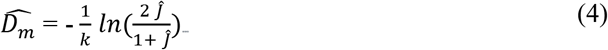

and

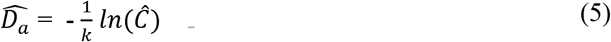

respectively; both the mash- and aaf-distances estimate the mutation distance between *A* and *B* [10, 21].

#### 4.2.3 Statistics properties of kssd

Due to the nature of Fisher-Yates shuffle [20], subspace ***s*** and sketch *S*(*A*) are an unbiased sampling (without replacement) of space ***S*** and the recoded *k*-mer set *r*(*A*), respectively; and we have |*r*(*A*)| = |*A*|, since the *k*-mer recoding function *r* is injective. Thus, we have:

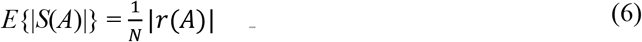

namely, the expected rate of dimensionality-reduction is 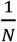 for sketch *S*(*A*).

For two *k*-mer sets *A* and *B*, we have:

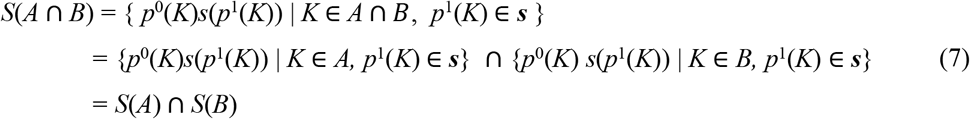

and similarly, we have:

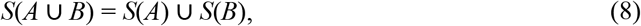

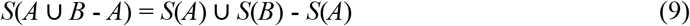

and

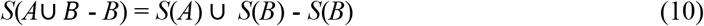

namely, the orders of kssd sketching operation and set operations are interchangeable. Since (*A ∩ B*) ∩ (*A ∪ B* – *A ∩ B*) = *ø*, we have |*S*(*A ∩ B*)| + |*S*(*A ∪ B* – *A ∩ B*)| = |*S*(*A ∪ B*)|, where |*S*(*A ∩ B*)| and |*S*(*A ∪ B* – *A* ∩ *B*)| are independent. Therefore, we have:

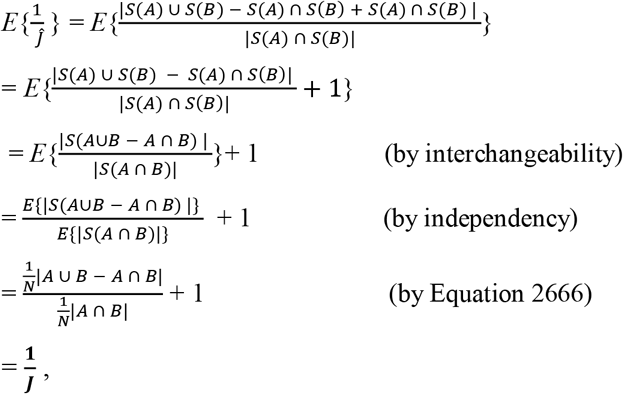

namely,

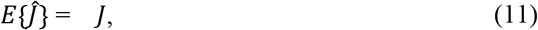

similarly, we have:

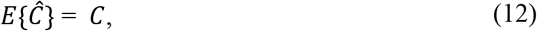

which means *Ĵ* and *Ĉ* are unbiased estimates of *J* and *C*, respectively. Both *Ĵ* and *Ĉ* are essentially sample proportions, thus have asymptotical gaussian distribution 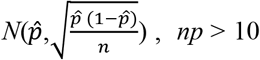, where 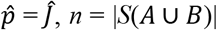 or 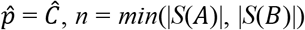 when *p* = *J* or *C*, respectively. Therefore, the population standard deviation is given by

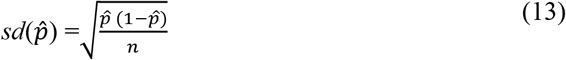

and the 95% confidence interval (CI) is 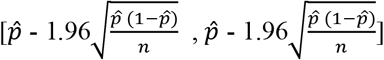. The *P*-value is defined by:

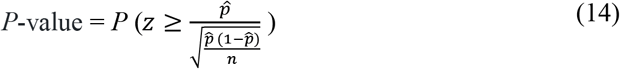

Where the random variable *z* follows the standardized normal distribution. To account for multiple testing problem, *Q*-value *Q* is obtained by multiplying the *P*-value *P* by the total number of comparisons, for example, for *x* queries search against *y* references, *Q* = *Pxy*.

#### 4.2.4 *k*-mer set decomposition

By applying Equation 1 532with subspace ***s_1_, s_2_***, …, ***s_n_*** individually, *k*-mer set *A* can be decomposed into a collection of sketches *d*(*A*) = {*S_1_*(*A*), *S_2_*(*A*), …, *S_N_*(*A*)}, respectively. Since {***s_1_, s_2_***,…, ***s_N_***} is a partition of *k*-mer substring space ***S***, we have 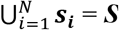 and ***s_i_*** ∩ ***s_j_*** = Ø, for 1 ≤ *i* ≤ *N* and 1 ≤ *j* ≤ *N*, therefore,

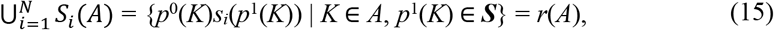

and

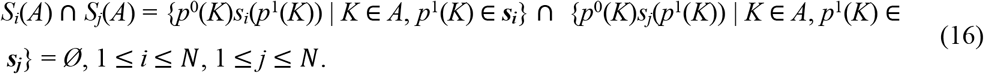

Thus, we have:

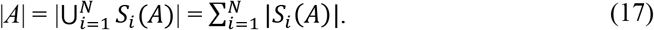

The above conversion from *A* to *d*(*A*) is termed as *A* decomposition and sketches *S_1_*(*A*) … *S_N_*(*A*) are termed as the components of *A*. Based on Equations 7–8,17 the ground truth Jaccard-coefficient *J* and containment-coefficient *C* of two *k*-mer sets *A* and *B* could be recovered by:

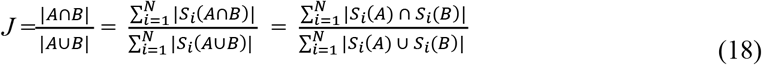

and

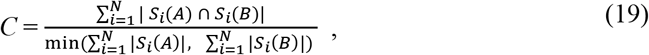

respectively. Equation 18–19 allow estimating the intersection and union of *A* and *B* component by component, where each component take only 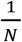 memory of the *k*-mer set, which is particularly useful when ground truth *J* and *C* is needed in a memory-limited computer.

#### 4.2.5 *k*-mer length

The optimal *k*-mer length for alignment-free sequences comparisons is roughly (but not strictly) framed by the Equation S2 in our previous work [22], namely:

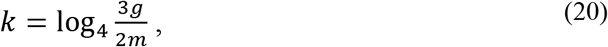

where *g* is the genome-size, and *m* is the allowed upper bound of the probability of random *k*-mer hitting (two *k*-mers are identical due to coincidence of random combination) which typically ranges from 0.001 to 0.01. Equation 20 is proposed under the considerations that *k* should be long enough to ensure *k*-mer uniqueness, while should be as short as possible to hold sensitivity [22]. It yields *k* = 16 for bacterial or smaller genomes, *k* = 20 or 22 for metagenomics/mammals’ or larger genomes and *k* = 18 for other genomes in-between. There is no necessary to use *k* > 22 and kssd avoids to do so to prevent *k*-mer substring space overflow (since the size of *k*-mer substring space is proportional to that of *k*-mer space by default).

#### 4.2.6 Implementation

*k*-mer substring space sampling/shuffling is implemented by the subcommand shuffle, for example:

~~~
kssd shuffle -k 9 -s 6 -l 3 -o out
~~~

this command will generate a file named ‘out.shuf’ which keeps the shuffled *k*-mer substring space, this file would then took as input for sequences sketching or decomposition. In current version kssd 1.0, users have no need to set kssp by himself, since kssd using an internal symmetric kssp ‘(0)_*n*_(1)_2*m*_(0)_*n*_’, where *n* and *2m* are the lengths of 0 or 1, respectively. Users need only to choose half-length of *k* (here set by -k 9) and half-length of *k*-mer substring (*m*, here set by `-s 6`). Such a design simplifies the *k*-mer substring selection step. The rate of dimensionality-reduction is controlled by -*l*, for this example which uses -*l* 3 will have an expected rate of dimensionality-reduction rate 16^3^. For decomposition purpose set -*l* 0.

To sketch/decompose reference sequences, just run:

~~~
kssd dist -r <reference files dir> -L out.shuf -o <ref_outdir>
~~~

or

~~~
kssd dist -r <reference files dir> -L 3 -k 9 -o <ref_outdir>
~~~

either command will generate a database of reference-sketches in folder `ref_outdir/ref`; the latter one actually combines the two commands below internally:

~~~
kssd shuffle -k 9 -s 6 -l 3 -o <ref_outdir>/default.shuf &&
kssd dist -r <reference files dir> -L ref_outdir/default.shuf -o <ref_outdir>
~~~

it simplifies the process at the cost of losing control for option ‘`-s`’.

To sketch/decompose query sequences, make sure using the same ‘.shuf’ file with the references, and just run:

~~~
kssd dist -o <qry_outdir> -L out.shuf|<ref_outdir>/default.shuf <query files dir>
~~~

it will generate a database of query-sketches in folder `qry_outdir/qry`. Then we can perform all queries versus all references comparisons by:

~~~
kssd dist -r <ref_outdir>/ref -o <outdir> <qry_outdir>/qry
~~~

To compare references to themselves, just run:

~~~
kssd dist -r <ref_outdir>/ref -o <outdir> <ref_outdir>/qry
~~~

then the distance will output to the ‘distance’ file in the folder `<outdir>`.

## 5 Supplementary Files

### 1. Resemblance Accuracy Experiment

The testing data sets, script, the shuffled *k*-mer substring space file (.shuf file), the detailed workflow (README.md) to generate Figure 1 and the source data of Figure 1.

### 2. Containment Accuracy Experiment

The testing data sets, the shuffled *k*-mer substring space file, sketches, the detailed workflow to generate Figure 2 and the source data of Figure 2.

## 6 Acknowledgments

This study was supported by National Key R&D Program of China (2018YFC1004500), the National Science Foundation of China (81872330, 31741077), and Basic Research Grant from Science and Technology Innovation Commission of Shenzhen Municipal Government (JCYJ20170817111841427).

